# Exfoliating Bark Does Not Protect *Platanus occidentalis* L. From Root-Climbing Lianas

**DOI:** 10.1101/046102

**Authors:** James R. Milks, J. Hibbard, Thomas P. Rooney

## Abstract

Lianas are structural parasites that depress growth, fertility and survival rates of their hosts, but the magnitude to which they alter these rates differ among host species. We tested the hypothesis that sycamore (*Platanus occidentalis* L.) would have fewer adventitious root-climbing lianas. We reasoned that because *P. occidentalis* possesses exfoliating bark, it would periodically shed newly-established lianas from the trunk. We investigated the distribution of lianas on the trunks of trees ≥10 cm DBH in floodplains in southwestern Ohio. Contrary to predictions, *P. occidentalis* trees had significantly more root-climbing lianas than expected at three of five sites, and significantly fewer than expected at one site. In contrast, members of the *Acer* genus (boxelder (*A. negundo* L.), sugar maple (*A. saccharum* L.) and silver maple (*A. saccharinum* L.) had less than half of the root-climbing lianas as expected. We find no support for our hypothesis that bark exfoliation protects *P. occidentalis* trees from root-climbing lianas in our study, and suggest possible mechanisms that might protect *Acer* species from adventitious root-climbing lianas.

## Introduction

Lianas are structural parasites that depress growth, fertility and survival rates of their hosts, but the magnitude to which they alter these rates differs among host species (Givnish 1992, 1995, Stevens 1987, Ladwig and Meiners 2009, van der Heijden and Phillips 2009, Ingwell et al. 2010). Different tree species exhibit different mechanisms to decrease the number of lianas successfully establishing. Proposed mechanisms include: compound leaves, fragile spines, and ant guards, flexible trunks, long leaves and high relative growth rates (Putz 1980, 1984, Givnish 1995).

Few studies have examined the role of bark shedding as a defense against lianas (e.g. Talley et al. 1996a; Carsten et al. 2002) and these have been confined to species in the tropics. Bark shedding would be expected to protect against liana infestation, as lianas would be expected to be shed along with pieces of bark. This should be especially effective against root-climbing lianas, as species with this growth form attach to bark to climb. Talley et al. (1996a) noted that bark shedding reduced lianas in two species of Australian rainforest trees. Carsten et al. (2002) found a more complex pattern, as liana densities increased at intermediate levels of bark shedding but decreased at higher levels of shedding.

Temperate floodplains in the eastern United States are well suited for studying liana/host relationships. Floodplain forests are subject to several factors that increase liana abundance, including disturbance through periodic flooding (van der Heijden and Philips 2008) and forest fragmentation (Londré and Schnitzer 2006). Floodplains are also the primary habitat of *Platanus occidentalis* L., a bark-shedding deciduous tree in the eastern United States (Burns and Honkala 1990). While bark-shedding has been hypothesized to protect *P. occidentalis* L. from lianas (Givnish 1992, 1995), no previous studies to our knowledge have tested this hypothesis.

Here, we tested the hypothesis that a temperate zone bark-shedding tree, *P. occidentalis* L., would have fewer root-climbing lianas than co-occurring species that do not shed bark. We counted the number of root-climbing lianas on tree trunks in five floodplain forests in southwestern Ohio. We predicted that *Platanus occidentalis* L. would have fewer lianas than expected compared to non bark-shedding species.

## Field-site Description

This study was conducted in mature floodplain forests at five different parks in the southwestern Ohio (39°30’ N, 84°0’ W): Germantown, Huffman Dam, Sugarcreek and Taylorsville Metroparks in Montgomery County, and The Narrows Preserve in Greene County. Montgomery County Parks within the Great Miami River Watershed, while The Narrows Preserve lies within the Little Miami River Watershed. Land use in both watersheds is predominantly cultivated cropland. Forest cover, pasture, and urban development are also present. Both watersheds are located within the Till Plains region of Ohio. This glaciated landscape contains rolling hills, moraines, and outwash plains (Zimmerman and Runkle 2010).

Floodplain forests are comprised of mature deciduous species. *Platanus occidentalis* L., *Acer negundo* L., *Celtis occidentalis* L., and *Populus deltoides* W. Bartram ex Marshall were the dominant species at our study sites. The invasive shrub *Lonicera maackii* (Rupr.) Herder is common in the forest shrub layer (Hutchinson and Vankat 1998).

## Methods

We recorded the diameter at breast height (DBH) and species of each tree ≥ 10 cm DBH in a single 10 m x 300 m belt transects (total 0.3 ha) in mature floodplain forests at five different parks. Each park was considered a separate site. Transects were randomly placed within the forests but in most cases, transects were within 50 meters of forest edges due to the narrow dimensions and fragmented nature of the floodplain forests in this region.

We tallied the number of adventitious root-climbing lianas present on the trunk of each tree at 1.6 m above ground level at the same time as the DBH measurements. Adventitious root-climbing lianas were chosen as they should be susceptible to being shed by trees with exfoliating bark. Data was collected in the spring over two field seasons (2007-2008).

We generated mean (± SE) lianas per tree, importance values and expected numbers of lianas per tree species for each site. For the purposes of our analyses, we combined *Fraxinus americana* and *Fraxinus pennsylvanica* into *Fraxinus* sp., as the two species are virtually indistinguishable in the field in our area. Importance values for each tree species were calculated by adding relative DBH and relative densities for each species, then dividing by 2 and multiplying by 100. We calculated relative DBH by dividing the total DBH for each species by the total DBH for all trees at the site.

Relative density was calculated totaling all individual stems per species and dividing by the total individual stems per site. Relative DBH was used as we expected larger trees to host more lianas than smaller trees due to increased age and having more surface area to which lianas could attach (Talley et al. 1996a, Buron et al. 1998, Carsten et al. 2002, Reddy and Parthasarathy 2006, Leicht-Young, et al. 2010). To test if this pattern held for our sites, we analyzed the number of lianas per tree versus DBH with a hurdle model for zero-inflated poisson distributions, using the pscl package for R (Zeileis et al. 2008, Jackson 2011, R Development Core Team 2011).

After finding a significant relationship between number of lianas and DBH, we examined differences in number of lianas per cm DBH per tree species and site using a two-way ANOVA with a post-hoc Tukey Honestly Significant Difference test. Finding a significant interaction term between site and species, we then analyzed differences between observed and expected lianas per tree species with replicated goodness of fit (G) tests, with each site as a replicate. We obtained expected numbers of lianas for each site and for each tree species by multiplying importance value for each tree species by the total number of lianas counted at each site.

## Results

We measured 1541 trees comprising 18 species and counted 1967 root-climbing lianas (mostly *Toxicodendron radcans* (L.) Kuntze and a few *Parthenocissus quinquefolia* (L.) Planch.) in a total of 1.5 ha (Table 1). Of those 18 tree species, *P. occidentalis* L. and *Acer negundo* L. had the highest importance values at 32.3 and 23.2, respectively. Larger trees were significantly more likely to have at least one liana than smaller trees (β_1_ = 0.025 ± 0.003 SE, P < 0.0001, Fig. 1a) and the number of lianas per tree also increased with DBH (β_1_ = 0.018 ± 0.001 SE, P < 0.0001, Fig. 1b). Sites differed greatly in liana abundance, from a low of 0.0042 ± 0.008 SE lianas per cm DBH per tree at Germantown to a high of 0.059 ± 0.004 SE lianas per cm DBH per tree at Huffman (F = 36.3, P <0.0001, Table 2, Fig. 2). There was a significant interaction term between site and species (F = 4.91, P <0.0001, Table 2), which was then examined using G-tests.

**Fig. 1.**
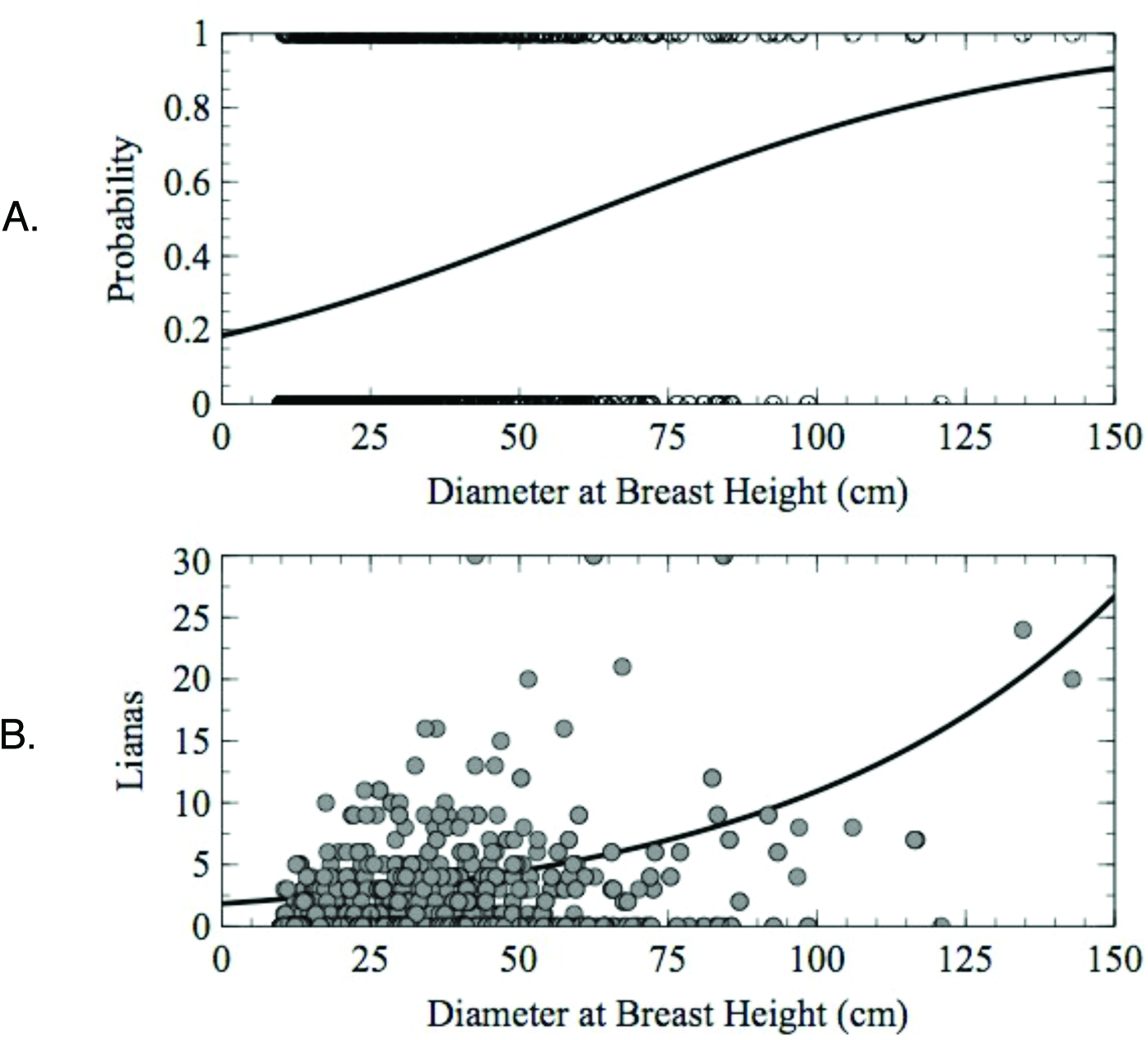
**A.** Estimated probability of at least one liana growing on a tree versus diameter at breast height along with the estimated logistic regression curve. **B**. Truncated Poisson regression of the number of lianas growing on trees versus diameter at breast height.

**Fig. 2.**
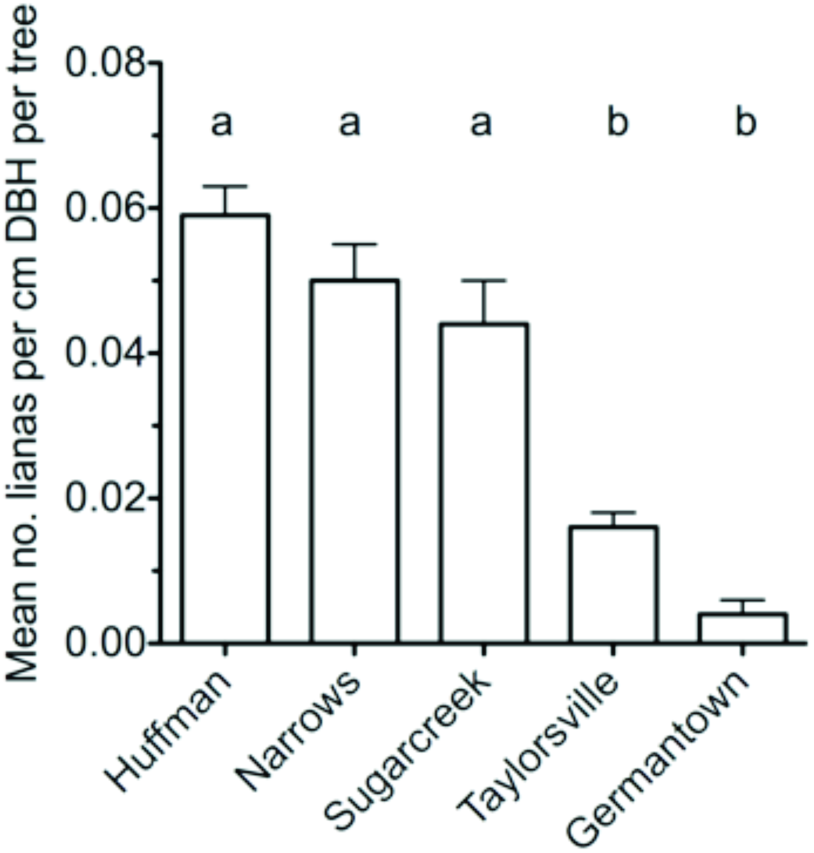
Mean (± SE) lianas per cm DBH per tree for each site. Sites that with significantly different means (p ≤ 0.05) based on Tukey’s HSD test are denoted with different letters.

**Table 1.** Species of trees, sites the species occurred at (sites), number of individual trees (n), number of lianas found on each species of tree, percentage of trees with at least one root-climbing liana clinging to them, and importance values per tree species for all sites combined. Sites are as follows: G = Germantown, H = Huffman Dam, N = The Narrows, S = Sugarcreek, and T = Taylorsville.

| Species | Sites | n | No. root-climbing lianas | Percent of trees Infested | Importance Value |
| --- | --- | --- | --- | --- | --- |
| *Acer negundo* | G, H, N, S, T | 432 | 169 | 14.1 | 23.2 |
| *Acer saccharum* | N | 30 | 4 | 13.3 | 1.4 |
| *Acer saccharinum* | G, H, T | 88 | 10 | 2.3 | 5.8 |
| *Aesculus glabra* | G, H, N, S, T | 88 | 67 | 37.5 | 4.7 |
| *Celtis occidentalis* | G, H, N, S, T | 163 | 482 | 71.8 | 10.1 |
| *Crataegus sp.* | H | 2 | 2 | 100 | 0.1 |
| *Fraxinus sp.* | H, N, S, T | 65 | 85 | 21.5 | 3.7 |
| *Juglans nigra* | G, H, N, S, T | 25 | 51 | 36.0 | 1.9 |
| *Liriodendron tulipifera* | S | 26 | 11 | 23.1 | 1.5 |
| *Maclura pomifera* | H, T | 28 | 56 | 64.3 | 1.8 |
| *Morus alba* | H | 2 | 0 | 0 | 0.1 |
| *Platanus occidentalis* | G, H, N, S, T | 405 | 735 | 42.2 | 32.3 |
| *Populus deltoides* | H, T | 71 | 167 | 39.4 | 7.1 |
| *Prunus serotina* | S | 8 | 13 | 37.5 | 0.4 |
| *Robinia psuedoacacia* | T | 2 | 8 | 100 | 0.1 |
| *Tilia americana* | H | 4 | 0 | 0 | 0.3 |
| *Ulmus americana* | G, H, N | 60 | 43 | 30.0 | 3.1 |
| *Ulmus rubra* | H, T | 42 | 64 | 42.9 | 2.3 |
| **Total** |  | **1541** | **1967** | **32.8** | **99.9** |

**Table 2.**
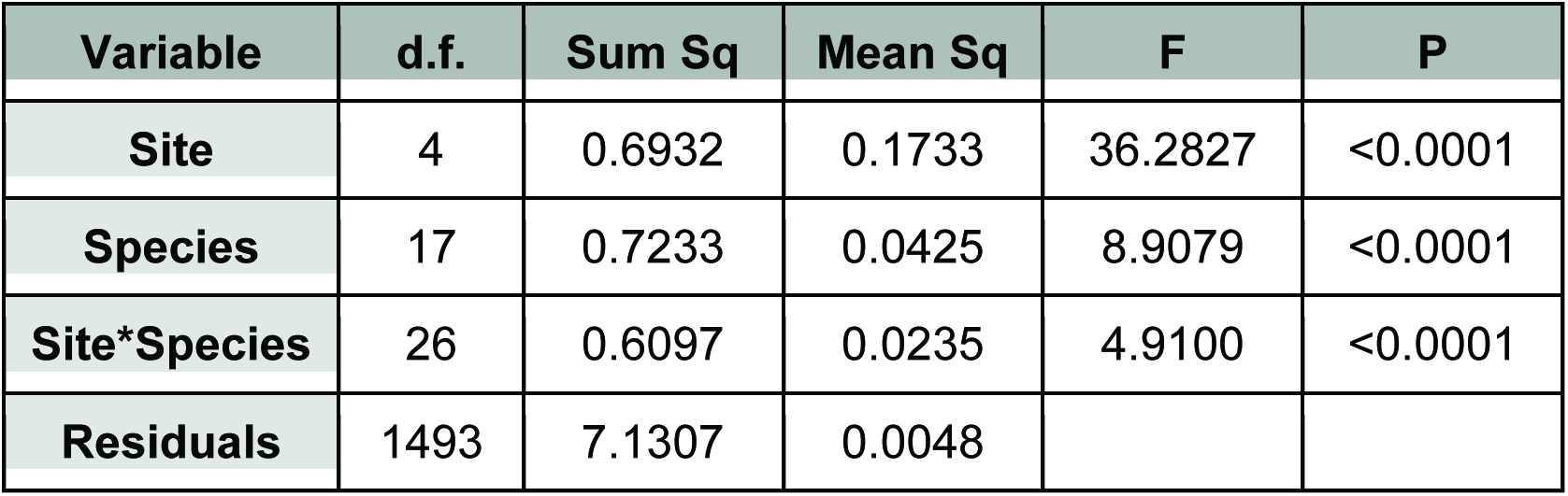
Variation in the number of lianas per cm DBH attributable to tree species, site and their interaction.

We found that the number of root-climbing lianas growing on *P. occidentalis* was significantly greater than expected at three of five sites, significantly less at one site and did not differ from expected abundance at the remaining site (Table 3). When data were pooled across sites, *P. occidentalis* had 30% more root-climbing lianas than expected (pooled G = 73.6, d.f. = 1, P <0.001). In contrast, *A. negundo* had significantly fewer lianas than expected at three sites and did not differ from expected abundance at two sites (Table 3). When data were pooled across sites, *A. negundo* had 59% fewer root-climbing lianas than expected (pooled G = 222.2, d.f. = 1, P <0.0001). *Acer saccharinum* had 76.5% fewer lianas at all three sites where individuals of this species occurred (pooled G = 32.8, d.f. = 1, P <0.0001; Table 3). A third maple species, *Acer saccharum,* had 83% fewer lianas than expected at the one site we found it (G = 45.8, d.f. = 1, P < 0.0001). Other species exhibited idiosyncratic relationships between site and liana abundance. *Fraxinus* spp. had significantly more lianas than expected at one site (G = 45.3, d.f. = 1, P < 0.0001) but fewer than expected at two other sites (G = 31.8, d.f. = 1, P < 0.0001 and G = 13.7, d.f. = 1, P < 0.0002; Table 3). When combined across all sites, there was no significant relationship (pooled G = 0.38, d.f. = 1, P = 0.54). A fifth species (*Celtis occidentalis* L.) had significantly more lianas than expected at two sites (G = 131.50, d.f. =1, P < 0.0001 and G = 33.96, d.f. = 1, P < 0.0001), significantly fewer than expected at two sites (G = 5.81, d.f. = 1, P = 0.016 and G = 10.4, d.f. = 1, P = 0.0013), and no significant difference at the fifth (G = 3.00, d.f. = 1, P = 0.083, Table 3). When combined across sites, *C. occidentalis* had 61% more vines than expected (pooled G = 114.7, d.f. =1, P < 0.0001).

**Table 3.** Number of stems, importance values per species per site, observed lianas abundance, expected liana abundance, and p-values for the five species of trees with overall importance values > 0.05 and which were found in three or more sites. N/A = species not recorded at that site. Bold P-value indicates statistical significance.

| Species | Site | No. stems | Importance value per site | Lianas Obs. | Lianas Exp. | P-value |
| --- | --- | --- | --- | --- | --- | --- |
| *Acer negundo* | Germantown | 120 | 58.4 | 11 | 13.43 | 0.3078 |
|  | Huffman | 142 | 22.9 | 59 | 244.46 | **<0.0001** |
|  | Narrows | 88 | 24.7 | 75 | 110.98 | **<0.0001** |
|  | Sugarcreek | 6 | 4.4 | 0 | 8.36 | **<0.0001** |
|  | Taylorsville | 76 | 14.3 | 24 | 36.51 | **0.0181** |
| *Acer saccharinum* | Germantown | 26 | 15.12 | 0 | 3.49 | **0.0059** |
|  | Huffman | 2 | 0.4 | 0 | 4.72 | **0.0021** |
|  | Narrows | 0 | 0.0 | N/A | N/A | N/A |
|  | Sugarcreek | 0 | 0.0 | N/A | N/A | N/A |
|  | Taylorsville | 60 | 13.4 | 10 | 34.28 | **<0.0001** |
| *Celtis occidentalis* | Germantown | 4 | 1.9 | 2 | 0.45 | 0.0831 |
|  | Huffman | 137 | 26.3 | 454 | 280.57 | **<0.0001** |
|  | Narrows | 6 | 1.7 | 2 | 7.52 | **0.0159** |
|  | Sugarcreek | 4 | 3.0 | 0 | 5.12 | **0.0013** |
|  | Taylorsville | 12 | 2.2 | 24 | 5.67 | **<0.0001** |
| *Fraxinus sp.* | Germantown | 0 | N/A | N/A | N/A | N/A |
|  | Huffman | 5 | 1.1 | 42 | 12.15 | **<0.0001** |
|  | Narrows | 29 | 9.2 | 39 | 41.47 | 0.6850 |
|  | Sugarcreek | 19 | 1.3 | 4 | 19.16 | **<0.0001** |
|  | Taylorsville | 12 | 2.7 | 0 | 6.78 | **0.0002** |
| *Platanus occidentalis* | Germantown | 24 | 18.3 | 5 | 4.21 | 0.6779 |
|  | Huffman | 72 | 20.8 | 286 | 221.71 | **<0.0001** |
|  | Narrows | 105 | 39.9 | 269 | 179.31 | **<0.0001** |
|  | Sugarcreek | 50 | 48.8 | 166 | 82.89 | **<0.0001** |
|  | Taylorsville | 154 | 41.8 | 48 | 106.87 | **<0.0001** |

## Discussion

We find no support from our data for the hypothesis that bark shedding protects *P. occidentalis* from root-climbing lianas. *P. occidentalis* had either the same as or more than the expected number of lianas at four sites out of five sites whereas we predicted that *P. occidentalis* should have fewer than expected lianas. This contrasts with Talley et al. (1996a) who found that bark-shedding trees in Queensland tropical forests had fewer than expected root-climbing lianas. Carsten et al. (2002) found that root-climbing lianas increased on trees with intermediate bark roughness and levels of bark-shedding and decreased at high levels of shedding and on trees with smooth bark. It is possible that *P. occidentalis* would be fall within the intermediate range of the bark texture scale of Carsten et al. (2002). One possible test would be to compare individual *P. occidentalis* for differences in bark shedding levels and liana load as individual *P. occidentalis* vary in levels of bark shedding with some trees shedding nearly all bark and others shedding very little (Milks, personal observation).

In contrast to *P. occidentalis, A. negundo, A. saccharum,* and *A. saccharinum* had either the expected number of lianas or significantly fewer lianas than expected on all sites where those species occurred. Other studies have also noted fewer than expected lianas on *A. saccharum.* Both Talley et al. (1996b) and Leicht-Young et al. (2010) found fewer than expected *T. radicans* lianas on *A. saccharum* in forests in Alabama, Indiana and Michigan. However, our finding of fewer than expected lianas on

A. *saccharinum* contrasts with Leicht-Young et al.’s (2010) finding. Only 2.3% of *A. saccharinum* in our study were infested with lianas whereas Leicht-Young et al. found an infestation rate of 28.6%. The reasons for this difference likely lie in differences in our methods, as we focused on root-climbing lianas whereas Leicht-Young et al. (2010) included all lianas, regardless of climbing mode. Another possible reason may be due to differences in the dominant liana species. Our dominant root-climbing species was *T. radicans* whereas *P. quinquefolia* dominated the liana community in Leicht-Young et al.‘s (2010) study.

Other tree species in our study (like *Fraxinus* sp. and *C. occidentalis*) showed no consistent trend between site and liana abundance, with some sites having more than expected lianas, others having fewer than expected. Site differences are apparent in our data, especially between Germantown and the other sites (Fig. 1). The reasons for those large differences in liana abundance between sites are not clear and may simply represent spatial heterogeneity.

Possible reasons for the differences between liana abundance between *P. occidentalis* and *A. negundo, A. saccharum* and *A. saccharinum* include leaf size, bark morphology and bark chemistry. Putz (1984) found that larger leaf size protected trees from lianas on Barro Colorado Island. However, in our area, *P. occidentalis* generally has larger leaves than any of the *Acer* species, making leaf size an unlikely mechanism in eastern temperate floodplain forests.

Bark morphology (smooth versus furrowed) is also unlikely to be an important mechanism, as this has been tested in other forest types with mixed results (Boom and Mori 1982, Carsten et al. 2002). In our study, *A. saccharum* and *A. negundo* had slightly furrowed bark whereas *A. saccharinum* had rough, furrowed bark. None of these three had many lianas, as only 14.1% of *A. negundo,* 13.3% of *A. saccharum* and 2.3% of *A. saccharinum* had lianas (Table 1). Bark morphology by itself is unlikely to explain our results, although it warrants further study.

One unexplored possibility is that allelopathic chemicals in the bark of some maple species may protect them from root-climbing lianas. Talley et al. (1996b) found allelopathic chemicals in *A. saccharum* bark (as well as chemicals in the bark of other tree species) could inhibit liana seedling germination and growth in the southern US, with differences in the presence of allelopathic chemicals influencing liana distributions on host trees. Talley et al. (1996a) found similar patterns in Australia. It is possible that bark chemistry may also protect *Acer* species from clinging lianas in our study region, although our study did not investigate that possibility.

This study is, to our knowledge, the first study that demonstrates that bark shedding in *P. occidentalis* does not protect that species from liana infestation. We also showed that *A. negundo* has either the same or fewer than expected lianas, which is also a new finding. Further research into the characteristics that decrease root-climbing liana abundance on *A. negundo* is desirable. Future investigations could also examine host preferences for different species of lianas in temperate floodplains, and whether variability in bark shedding among individual *P. occidentalis* individuals affects liana loads.

## Acknowledgements

The idea for this project was first suggested by T. Givnish in his class entitled The Vegetation of Wisconsin.

